# DeepViral: infectious disease phenotypes improve prediction of novel virus–host interactions

**DOI:** 10.1101/2020.04.22.055095

**Authors:** Wang Liu-Wei, Şenay Kafkas, Jun Chen, Nicholas Dimonaco, Jesper Tegnér, Robert Hoehndorf

## Abstract

**Motivation:** Infectious diseases from novel viruses have become a major public health concern. Rapid identification of virus–host interactions can reveal mechanistic insights into infectious diseases and shed light on potential treatments. Current computational prediction methods for novel viruses are based mainly on protein sequences. However, it is not clear to what extent other important features, such as the symptoms caused by the viruses, could contribute to a predictor. Disease phenotypes (i.e., signs and symptoms) are readily accessible from clinical diagnosis and we hypothesize that they may act as a potential proxy and an additional source of information for the underlying molecular interactions between the pathogens and hosts.

**Results:** We developed DeepViral, a deep learning based method that predicts protein–protein interactions (PPI) between humans and viruses. Motivated by the potential utility of infectious disease phenotypes, we first embedded human proteins and viruses in a shared space using their associated phenotypes and functions, supported by formalized background knowledge from biomedical ontologies. By jointly learning from protein sequences and phenotype features, DeepViral significantly improves over existing sequence-based methods for intra- and inter-species PPI prediction. Lastly, we propose a novel experimental setup to realistically evaluate prediction methods for novel viruses.

**Availability:** https://github.com/bio-ontology-research-group/DeepViral

## 1 Introduction

Infectious diseases emerging unexpectedly from novel and reemerging pathogens have been a major enduring public health concern around the globe (Jones *et al*., 2008). Pathogens disrupt host cell functions (Finlay and Cossart, 1997) and target immune pathways (Dyer *et al*., 2010) through complex inter-species interactions of proteins (Dyer *et al*., 2008), RNA (Fajardo *et al*., 2015) and DNA (Weitzman *et al*., 2004). The study of pathogen–host interactions (PHI) can therefore provide insights into the molecular mechanisms underlying infectious diseases and guide the discoveries of novel therapeutics or provide a basis for the repurposing of available drugs. For example, a previous study of many PHIs showed that pathogens typically interact with the protein hubs (those with many interaction partners) and bottlenecks (those of central locations to important pathways) in human protein–protein interaction (PPI) networks (Dyer *et al*., 2008). However, due to cost and time constraints, experimentally validated pairs of interacting pathogen–host proteins are limited in number. Therefore, the computational prediction of PHIs is a useful complementary approach in suggesting candidate interaction partners from the human proteome.

Existing PHI prediction methods for novel viruses typically utilize protein sequence features of the interacting proteins (Eid *et al*., 2016; Zhou *et al*., 2018; Alguwaizani *et al*., 2018; Yang *et al*., 2020). While protein functions have been shown to predict intra-species (e.g., human) PPIs (Guzzi *et al*., 2011; Jain and Bader, 2010; Pesquita *et al*., 2009) and such protein specific features exist for some extensively studied pathogens, such as *Mycobacterium tuberculosis* (Huo *et al*., 2015) and HIV (Mukhopadhyay *et al*., 2014), for most pathogens, these features are rare and expensive to obtain. As new virus species continue to be discovered (Woolhouse *et al*., 2012), a method is needed to rapidly identify candidate interactions from information that can be obtained quickly, such as the signs and symptoms of the host, which may be utilized as a proxy for the underlying molecular interactions between host and pathogen proteins.

The phenotypes elicited by pathogens, i.e., the signs and symptoms observed in a patient, may provide information about molecular mechanisms (Gkoutos *et al*., 2018). The information that phenotypes provide about molecular mechanisms is commonly exploited in computational studies of Mendelian disease mechanisms (Oellrich *et al*., 2016; Schofield *et al*., 2012), for example to suggest candidate genes (Hoehndorf *et al*., 2011; Meehan *et al*., 2017) or diagnose patients (Köhler *et al*., 2009), but the information can also be used to identify drug targets (Hoehndorf *et al*., 2013a) or gene functions (Hoehndorf *et al*., 2013b). We hypothesize that the host phenotypes elicited by an infection with a pathogen are, among others, the result of molecular interactions, and that knowledge of the phenotypes exhibited by the host can be used to suggest the protein perturbations from which these phenotypes arise.

One major challenge of the novel PHI prediction problem is the lack of ground truth negative data. A recent method, DeNovo (Eid *et al*., 2016), adopted a “dissimilarity-based negative sampling”: for each virus protein, the negatives are sampled from human proteins that do not have known interactions with other similar virus proteins (above a certain sequence similarity threshold). Another method based on protein sequences (Zhou *et al*., 2018), samples negatives from only the set of host proteins that are less than 80% similar (in terms of sequence similarity) to the host proteins in the positive training data. However, the influence of sequence similarity on function is not uniform and while there is evidence for a number of general evolutionary rules, we are unable to determine cutoffs for any specific protein or function (Whisstock and Lesk, 2003; Ponting, 2001). By construction, these sampling schemes make the human proteins in the negative set different from the positive set; when used not only for training a model but also for evaluating its performance, this sampling scheme has the potential to over-estimate the actual performance for finding novel PHIs. In a more realistic evaluation for a novel virus species, a model would be evaluated on all the host proteins with which it could potentially interact, regardless of sequence similarity.

From these motivations, we developed a machine learning method, DeepViral, to predict potential interactions between viruses and all human proteins for which we can generate the relevant features. Firstly, the features of phenotypes, functions and taxonomic classifications are embedded in a shared space for human proteins and viruses. We then extended a sequence model by incorporating the phenotype features of viruses into the model. We show that the joint model trained on both the sequences and phenotypes can significantly outperform state-of-the-art methods and predict potential PHIs in a realistic experimental setup for novel viruses.

## 2 Materials and methods

DeepViral is a model that predicts potential protein interactions between viruses and human hosts from the protein sequences and feature embeddings of phenotypes, functions and taxonomies. To enable predictions based on such different features we embedded them in a shared representation space. We then combine these feature embeddings with a protein sequence model to predict potential PHIs of novel viruses. The workflow of DeepViral is illustrated in Figure 1.

**Fig. 1:**
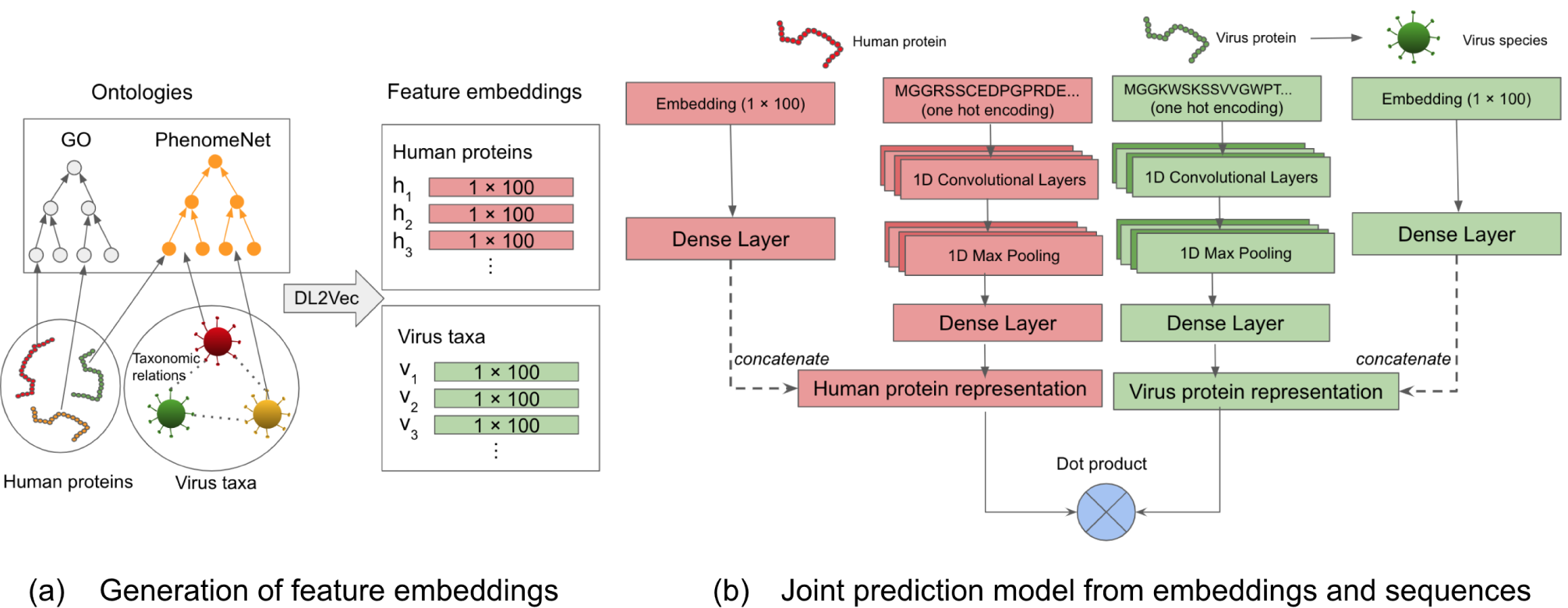
The workflow of DeepViral. (a) Generation of an embedding: the arrows of human proteins and virus taxa represent their annotations to the ontology classes. The dashed lines between viruses represent their taxonomic relations. The annotations, taxonomic relations and ontologies were fed into DL2Vec to generate feature embeddings of dimension 100 for each human protein and virus taxa. (b) Joint prediction model: latent representation learned from feature embeddings and protein sequences are concatenated into a joint representation, for human protein and virus protein respectively, on which a dot product is performed to predict interactions.

### 2.1 Data sources

Interactions between hosts and pathogens were obtained from the Host Pathogen Interaction Database (HPIDB; version 3) (Ammari *et al*., 2016). The phenotypes associated with pathogens were collected from the PathoPhenoDB (Kafkas *et al*., 2018), a database of manually curated and text-mined associations of pathogens, infectious diseases and phenotypes. We downloaded the PathoPhenoDB database version 1.2.1 (http://patho.phenomebrowser.net/).

The phenotypes associated with human genes were collected from the Human Phenotype Ontology (HPO) database (Köhler *et al*., 2018), and the phenotypes associated with mouse genes and the orthologous gene mappings from mouse genes to human genes originated from the Mouse Genome Informatics (MGI) database (Smith *et al*., 2018). The Entrez gene IDs in HPO and MGI were mapped to reviewed Uniprot protein IDs using the Uniprot Retrieve/ID mapping tool (https://www.uniprot.org/uploadlists) on March 6, 2020. The Gene Ontology annotations of human proteins (release date 2020-02-22) were downloaded from the Gene Ontology Consortium (Ashburner *et al*., 2000; The Gene Ontology Consortium, 2017). Human PPI networks were downloaded from String (Szklarczyk *et al*., 2019) and filtered to only include the interactions with experimental evidence. The human protein sequences were obtained from the Swiss-Prot database (Consortium, 2019).

To add background knowledge from biomedical ontologies of phenotypes and GO classes, we downloaded the cross-species PhenomeNET ontology (Hoehndorf *et al*., 2011; Rodríguez-García *et al*., 2017), from the AberOWL ontology repository (Hoehndorf *et al*., 2015a) on September 13, 2018. We obtained the NCBI Taxonomy classification (Sayers *et al*., 2009) as an ontology in OWL format (version 2018-07-27) from EMBL-EBI ontology repository (https://www.ebi.ac.uk/ols/ontologies/ncbitaxon).

### 2.2 Learning feature embeddings

To generate feature embeddings, we used DL2Vec (Chen *et al*., 2020), a recent method for learning features for entities (in our case, the human proteins and viruses) from their associations to ontological classes. DL2Vec first converted the ontologies and entity associations into a graph, with the classes and entities as the nodes and the associations and ontology axioms as the edges. Then a number of random walks were performed, starting from the entities over to the ontology graph and thereby generating a corpus of walks in the form of sentences capturing the graph neighborhoods and thereby the ontology axioms. After the construction of such sentences, a Word2vec skipgram model (Mikolov *et al*., 2013) was used to learn an embedding for each entity by learning from the corpus. Following the recommendations of the authors of DL2Vec, we fixed the number of walks to 100, the walk length to 30, the embedding dimension to 100 and the number of training epochs to 30. The embeddings were trained with the Word2Vec library in Julia (version 1.0.4). The resultant embedding was a vector representation of an entity capturing its co-occurrence relations with other entities within the walks generated by DL2Vec. As an example, the walks starting from a virus node explored its graph neighborhood, i.e., its associated phenotypes and its taxonomic relatives, and as an result, its feature embedding captured this information according to the co-occurrence patterns.

### 2.3 Supervised prediction models and parameter tuning

The neural network model of DeepViral consists of two components: a phenotype model based on the feature embeddings of viruses and human proteins and a sequence model based on the amino acid sequences of the human and viral proteins. The maximum input length of protein sequences is set to 1,000 amino acids and all shorter sequences are repeated up to the maximum length. The input protein sequences are encoded as a one-hot encoding matrix of 22 rows that represents each amino acid type and the original sequence length (before being repeated), and 1,000 columns representing each position of the amino acid sequence.

To predict the likelihood of an interaction between a pair of proteins, we trained the network as a binary classifier, to minimize the binary cross-entropy loss defined below,

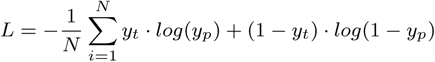

where *N* is the total number of predictions, *yt* and *yp* is the true label and predicted likelihood of *y*.

We implemented our model using the Keras library (Chollet *et al*., 2015) and performed training on Nvidia Tesla V100 GPUs. The phenotype model consists of a fully connected layer with the feature embeddings as input. The sequence model is a convolutional neural network (CNN) with the sequences as input and consists of 1-dimensional convolution, max pooling and fully connected layers. We tuned the following hyperparameters of the model: the sizes and numbers of the convolution filters, the size of the max pool and the number of neurons in the fully connected layers. We fixed these hyperparameters throughout all the experiments: 16 convolutional layers for each filter of 8, 16, …, 64 in length, a pool size of 200 and 8 neurons for the dense layers. We also used dropouts (Srivastava *et al*., 2014) for the convolutional and dense layers with a rate of 0.5 and LeakyReLU as the activation function for the dense layer with an alpha set to 0.1.

## 3 Results

### 3.1 Embedding features of viruses and human proteins from phenotypes, functions and taxonomies

We started with the biological hypothesis that phenotypes (i.e., symptoms) elicited by viruses in their hosts can act as a proxy for the underlying molecular mechanisms of the infection, and therefore may provide additional information to the prediction of potential PHIs for novel viruses.

To generate feature embeddings for human proteins and virus taxa, we applied a recent representation learning method DL2Vec (Chen *et al*., 2020), which learned feature embeddings for entities based on their annotations to ontology classes (see Section 2.2). DL2Vec takes two types of inputs: the associations of the entities with ontology classes (e.g., human proteins and their functions), and the ontologies themselves.

For representing virus taxa through the phenotypes they elicit in their hosts, we used the phenotype associations for viruses from PathoPhenoDB (Kafkas *et al*., 2018), a database of pathogen to host phenotype (signs and symptoms) associations. To increase the coverage of phenotypes beyond PathoPhenoDB, the taxonomic relations of the viruses were added from the NCBI Taxonomy (Sayers *et al*., 2009). By adding these taxonomic relations (as annotations of viruses to DL2Vec), we propagated the known phenotypes along the taxonomic hierarchies and learned a generalized embedding for viruses that do not have any phenotype annotations in PathoPhenoDB but have close relatives that do.

Similarly, for representing human proteins, we used the annotations of their associated phenotypes from the Human Phenotype Ontology (HPO) database (Köhler *et al*., 2018), the phenotypes associated with their mouse orthologs from the Mouse Genome Informatics (MGI) database (Smith *et al*., 2018), and their protein functions from the Gene Ontology (GO) database (Ashburner *et al*., 2000; The Gene Ontology Consortium, 2017). We propagated these annotations through the human PPI network, which has been shown to improve prediction for gene-disease associations (Alshahrani and Hoehndorf, 2018).

To provide DL2Vec with structured background knowledge of human and mouse phenotypes as well as protein functions, we used the cross-species phenotype ontology PhenomeNET (Hoehndorf *et al*., 2011; Rodríguez-García *et al*., 2017), which is built upon and includes the Gene Ontology (Ashburner *et al*., 2000; The Gene Ontology Consortium, 2017). These ontologies contain formalized biological background knowledge (Hoehndorf *et al*., 2015b), which has the potential to significantly improve the performance of these features in machine learning and predictive analyses (Smaili *et al*., 2019; Kulmanov *et al*., 2020).

### 3.2 A joint model for PPI prediction from sequences and phenotypes

DeepViral consists of a phenotype model trained on phenotypes caused by a viral infection and a sequence model trained on protein sequences, as shown in Figure 1 (b). The two models take a pair of virus and human proteins as input and predicts the probability score of their interaction. The inputs for a human protein are its feature embedding and its sequence, and the features for a viral protein are its sequence and the feature embedding of the virus species to which it belongs. The sequence model projects the protein sequence into a low dimension vector representation, which is concatenated with the vector projected from the embedding by the phenotype model to form a joint representation of the proteins. A dot product was performed over the two vector representations of the pair of proteins to compute their similarity, which was then used as input to a sigmoid activation function to compute their predicted probability of interaction. In an evaluation where the inputs were not symmetric, e.g., only using the feature embeddings of human proteins but not viruses (or vice versa), an additional dense layer was added to project the longer representation to the same dimension as the other so that the dot product could be performed.

Existing prediction methods for inter-species PPI (e.g., virus–human interactions) have rarely been compared with methods designed for intra-species (e.g. human) PPI prediction. To compare with the existing sequence-based methods for both intra- and inter-species PPI prediction, we evaluated DeepViral and RCNN (Chen *et al*., 2019), a recent method designed for intra-species prediction, on an existing dataset (Eid *et al*., 2016) that has been used to evaluate a number of PHI prediction methods (Yang *et al*., 2020; Alguwaizani *et al*., 2018; Zhou *et al*., 2018). The respective model performances and implementation details are shown in Supplementary Table 1. DeepViral trained only on sequences achieves comparable performance with other sequence based methods, while the joint model is able to achieve the best performances in most metrics. However, the evaluation dataset suffers from several drawbacks: 1) negative sampling (to create a balanced dataset) was based on sequence dissimilarity; 2) the training and test sets are small relative to the current size of the PHI databases; 3) there are overlapping viruses (i.e., data leakage) at species level between the training and test sets, which makes it unsuitable for the problem of novel PHI prediction.

**Table 1.**
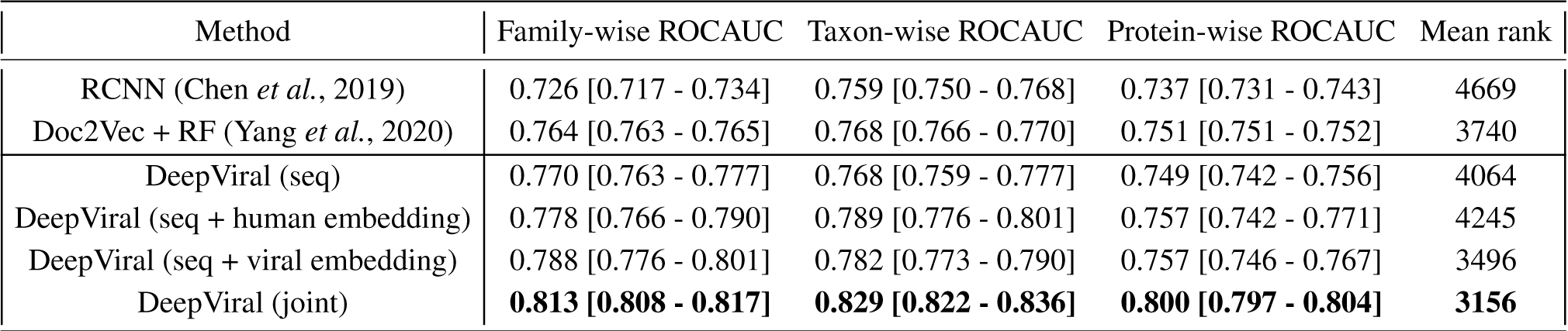
Comparison with the state-of-the-art methods on our dataset to evaluate the performances for novel viruses. The brackets after DeepViral indicate the features used for the model: seq – protein sequences, joint – both sequences and embeddings of human proteins and viruses. The square brackets behind ROCAUC scores indicate the 95% confidence interval.

| Method | Family-wise ROCAUC | Taxon-wise ROCAUC | Protein-wise ROCAUC | Mean rank |
| --- | --- | --- | --- | --- |
| RCNN (Chen *et al.*, 2019) | 0.726 [0.717 - 0.734] | 0.759 [0.750 - 0.768] | 0.737 [0.731 - 0.743] | 4669 |
| Doc2Vec + RF (Yang *et al.*, 2020) | 0.764 [0.763 - 0.765] | 0.768 [0.766 - 0.770] | 0.751 [0.751 - 0.752] | 3740 |
| DeepViral (seq) | 0.770 [0.763 - 0.777] | 0.768 [0.759 - 0.777] | 0.749 [0.742 - 0.756] | 4064 |
| DeepViral (seq + human embedding) | 0.778 [0.766 - 0.790] | 0.789 [0.776 - 0.801] | 0.757 [0.742 - 0.771] | 4245 |
| DeepViral (seq + viral embedding) | 0.788 [0.776 - 0.801] | 0.782 [0.773 - 0.790] | 0.757 [0.746 - 0.767] | 3496 |
| DeepViral (joint) | **0.813 [0.808 - 0.817]** | **0.829 [0.822 - 0.836]** | **0.800 [0.797 - 0.804]** | **3156** |

### 3.3 Experimental setup, negative sampling and evaluation metrics for novel viruses

Motivated by the need for more representative datasets to evaluate methods for novel PHI prediction, we constructed a larger dataset from the curated virus–host interactions in HPIDB (Ammari *et al*., 2016), a database of host–pathogen protein–protein interactions. We constructed our positive set by filtering HPIDB to include all virus–host interactions that 1) are provided with an MIscore, a confidence score for molecular interactions (Villaveces *et al*., 2015); 2) are associated with an existing virus family in the NCBI taxonomy (Sayers *et al*., 2009); 3) are within 1,000 amino acids in length (for both human and viral proteins). The sequence length cut-off of 1,000 is chosen to include over 88.2% of the human proteins in Swiss-Prot and over 91.6% of the virus proteins in HPIDB. After filtering, the dataset includes 24,678 positive interactions and 1,066 viral proteins from 14 virus families and 292 virus taxa.

To realistically evaluate the prediction performance, we performed a leave-one-family-out (LOFO) cross validation: at each run, one virus family in our positive set was left out for testing, 20% of the remaining families for validation, and the rest 80% for training. The objective of the LOFO cross-validation is to evaluate the model under a scenario in which the novel virus emerges from a novel virus family - in our study, “novel” is defined as the situation in which we have no or very little knowledge about its protein interactions and the molecular functions of the viral proteins.

Instead of using “dissimilarity-based negative sampling” to construct a balanced dataset, we sampled our negatives from all the possible pairwise combinations of human and viral proteins, as long as the pair did not occur in the positive set. Essentially, we treated all “unknown” interactions as negatives. As the dataset was at this point unbalanced with more negatives than positives, we evaluated the model with the area under the receiver operating characteristic (ROC) curve (Fawcett, 2006). A high ROCAUC indicates the ability of the model to prioritize the true positive interacting proteins out of all the human proteins. We computed a ROCAUC for each virus family, and also for each viral protein and virus taxon in that family, for which we reported the mean across them, i.e. macro averages. Each model was evaluated 5 times independently to compute the 95% confidence interval of the ROCAUC, which is bounded by 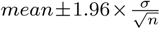, where *n* is the sample size and *σ* is the standard deviation. Additionally, the mean ranks of the true positive proteins were provided as a more interpretable metric: for each viral protein, we ranked all of the 16,627 human proteins in Swiss-Prot (with a length limit of 1,000) as its potential interaction partner based on the prediction score and obtained the mean ranks of the true positives.

### 3.4 Phenotypes improve prediction for novel viruses

With the newly constructed dataset, we further evaluated the existing methods as well as the variants of DeepViral, under the scenario in which a novel virus (from a novel family) emerges and no previous knowledge (except about its protein sequences and the phenotypes elicited in its hosts) is known.

We compared DeepViral with two existing state-of-the-art methods based on protein sequences: Doc2Vec + RF (Yang *et al*., 2020), a recent method predicting for virus–human interactions; and RCNN (Chen *et al*., 2019), a recent deep learning based method for intra-species (e.g., human) PPI prediction. To adapt Doc2Vec + RF on our dataset, we used the pretrained Doc2Vec model provided by the authors and the same parameters for the random forest model for training. Similarly, for RCNN, we used the pre-trained embeddings for amino acids and the same model parameters for training. Since the stop criterion for Doc2Vec + RF was to have at most 2 samples at each leaf node, we did not use validation data and trained it with the entirety of the training data, while a validation set was used for both RCNN and DeepViral as described in the experimental setup.

The performance of each model is shown in Table 1. For models using only sequence features, DeepViral and Doc2Vec + RF perform on a similar level across the metrics. As the current state-of-the-art method for intra-species PPI prediction, RCNN consistently yields the lowest performances. Adding human or virus embeddings individually shows a slight improvement in most metrics, compared to the sequence-only models, while the joint model with both embeddings achieved the best performances overall.

## 4 Discussion

We developed DeepViral, a machine learning method for predicting PHIs between viruses and human hosts. DeepViral is, based on our review of the literature, the first predictor using clinical phenotypes as an additional feature in PHI prediction and it has been seen to provide a significant improvement (*p <* 0.05; see confidence intervals in Table 1) over purely sequence based methods. Phenotype-based approaches have been successful in predicting disease-gene associations for Mendelian diseases (Hoehndorf *et al*., 2011) and intra-species PPIs (Alshahrani *et al*., 2017), but have not yet been used for the prediction of (inter-species) PHIs in infectious diseases. Our model avoids the bottleneck of identifying the molecular functions of pathogen proteins by instead introducing a novel and – in the context of infectious diseases – rarely explored type of feature, the phenotypes elicited by pathogens in their hosts, as a “proxy” for the molecular mechanisms, which in turn eventually produce the observed clinical phenotypes.

The focus of our method on utilizing features generated based on endo-phenotypes observed in humans and mice (Schofield *et al*., 2016) has therefore the crucial advantage that we can identify host-pathogen interactions that may contribute to particular signs and symptoms. For example, our model consistently prioritizes the interaction between the proteins of Zika virus (`NCBITaxon:64320`) and DDX3X (`UniProt:O00571`) in humans. Infections with Zika virus have the potential to result in abnormal embryogenesis and, specifically, microcephaly (Wang *et al*., 2017). Phenotypes associated with DDX3X in the mouse ortholog include abnormal embryogenesis, microcephaly, and abnormal neural tube closure (Chen *et al*., 2016). DDX3X mutations in humans have been found to result in intellectual disability, specifically in females and affect individuals in a dose-dependent manner (Blok *et al*., 2015). While DDX3X has previously been linked to the infectivity of the Zika virus (Doñate-Macián *et al*., 2018), our model further suggests a role of DDX3X in the development of the embryogenesis phenotypes resulting from Zika virus infections.

While we demonstrate quantitatively an improvement over existing methods on an existing dataset (Eid *et al*., 2016), we argue that the performances using this evaluation approach may have been over-estimated due to the negative sampling scheme based on sequence similarity that is used not only for training but also for evaluation of the model. Under a more realistic evaluation procedure that considers all host proteins as potential interaction partners for novel viruses, the achieved predictive performances are considerably lower. This calls for future efforts in the direction of PHI prediction of novel viruses, an issue today of increasing relevance to global public health. Accurate predictions of potential PHIs for novel pathogens with rapidly obtainable features would be an important development for understanding infectious disease mechanisms and the repurposing of existing drugs.

An example of such a novel virus is the novel coronavirus SARS-CoV-2, which as of 6th August 2020 reached more than 18 million infected cases and 707 thousand fatalities globally (Dong *et al*., 2020) in a timespan of 9 months. Based on a recently released dataset of 332 PHIs from 26 viral proteins of SARS-CoV-2 (Gordon *et al*., 2020), we applied DeepViral by treating it as a novel family (with no other Coronaviridae viruses in the dataset) and achieved a family-wise ROCAUC of 0.723 (0.699– 0.747; 95% CI), which is within the observed variability in predicting for different virus families, as shown in Figure 2. This family-wise variability suggests that the learned features to predict for PHIs may have different generalization power across families, possibly a result of varying degrees of (dis)similarity between the virus families. Nonetheless, optimizing the predictive power for a single virus, e.g., SARS-CoV-2, requires a case-by-case experimental setup. Specifically in the case of SARS-CoV-2, one can potentially relax the leave-one-family-out evaluation, as we have prior knowledge about other species in its family, e.g., SARS-CoV and MERS-CoV, and their interactions with hosts and protein functions (Thiel *et al*., 2003). This is indeed a topic for further investigation.

**Fig. 2:**
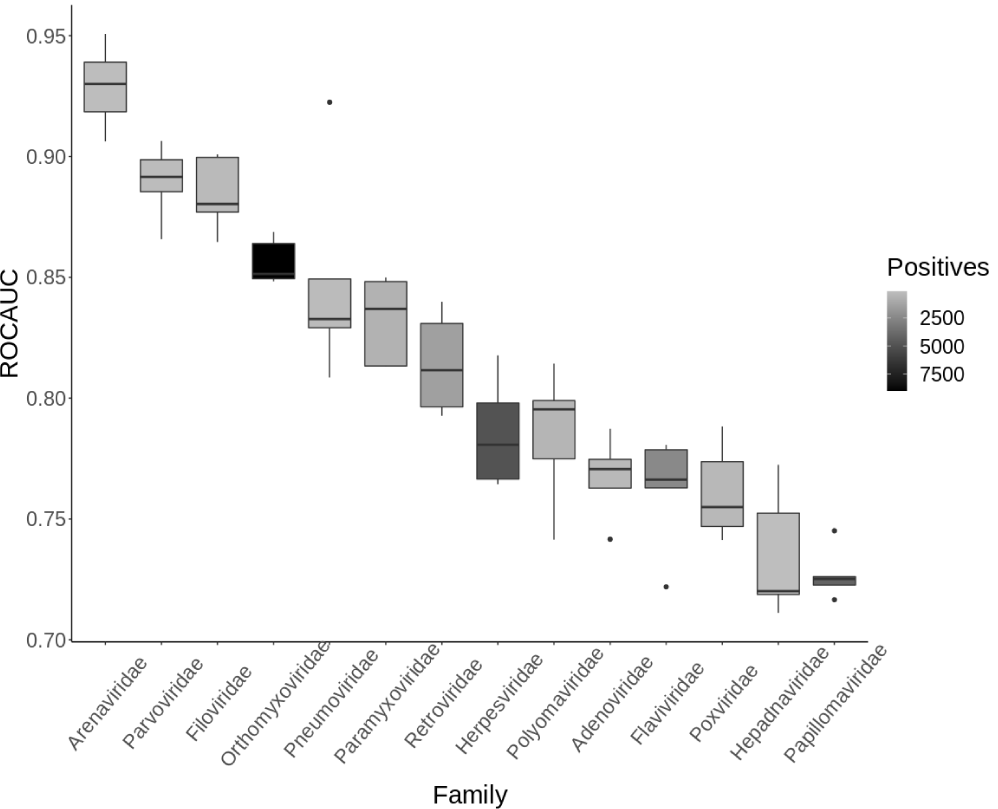
ROCAUC for each of the 14 virus families from the joint model, colored by the number of positives belonging to that family.

There are several limitations that can be addressed by future efforts. One is the scarcity of training data for inter-species PPIs and this may be leveraged by transfer learning on the much larger intra-species PPI data available for humans and other model organisms. We also ignored other types of PHIs outside virus–human interactions in our current study, such as those of other hosts, e.g., plants and fishes, and other types of pathogens, e.g., bacteria and fungi. Additionally, predicting tissue-specific PHIs would also provide additional insights, as proteins of both human hosts (Fagerberg *et al*., 2014) and viruses (Jarosinski *et al*., 2012) often have tissue-specific expressions and functions.

## Acknowledgements

We would like to thank Maxat Kulmanov and Mona Alshahrani for their advice on earlier versions of this work. We also thank Jeffery Law for making public the mappings of the SARS-CoV-2 proteins.

