## Supplementary Table 1 for "DeepViral: infectious disease phenotypes improve prediction of novel virus–host interactions"

### 1 Comparison with existing methods

| Method | SN (%) | SP (%) | ACC (%) | PPV (%) | NPV (%) | MCC | AUC | F1 (%) |
|---|---|---|---|---|---|---|---|---|
| DeNovo [Eid et al., 2016] | 80.71 | 83.06 | 81.90 | NA | NA | NA | NA | NA |
| VirusHostPPI [Zhou et al., 2018] | 80.00 | 88.94 | 84.47 | 87.86 | 81.64 | 0.692 | 0.897 | NA |
| Doc2Vec + RF [Yang et al., 2020] | **90.33** | 96.17 | 93.23 | 95.99 | 90.74 | 0.866 | **0.981** | 93.07 |
| RCNN [Chen et al., 2019] | 89.88 | 95.58 | 92.73 | 95.38 | 90.46 | 0.857 | 0.974 | 92.52 |
| DeepViral (seq) | 89.36 | 96.89 | 93.13 | 96.68 | 90.13 | 0.865 | 0.960 | 92.86 |
| DeepViral (seq + human embedding) | 88.43 | 96.22 | 92.32 | 95.94 | 89.23 | 0.849 | 0.955 | 92.02 |
| DeepViral (seq + viral embedding) | 88.29 | 97.24 | 92.76 | 96.97 | 89.26 | 0.859 | 0.967 | 92.42 |
| DeepViral (joint) | 90.27 | **97.58** | **93.91** | **97.43** | **90.93** | **0.881** | 0.976 | **93.68** |

Table 1: Comparison with the state-of-the-art methods on the datasets of [Eid et al., 2016] (the performances of first 3 methods are from the original papers respectively). RCNN and the variants of DeepViral are evaluated 5 times independently to compute the mean of the metrics: SN - sensitivity, SP - specificity, ACC - accuracy, PPV - positive predictive value (precision), NPV - negative predictive value, MCC - Matthews correlation coefficient, AUC - area under the ROC curve. DeepViral (seq) only utilizes the protein sequences and the joint model also includes both the human and virus embeddings as input. The bold numbers represent the best metric for a dataset.

**Implementation details:** The dataset contains contains 5,020 positives and 4,734 negatives in the training set, and 425 positives and 425 negatives in the testing set. Since no validation set was used previously, we constructed a validation set by randomly sampling 10% of the training set, which was used for choosing the best epoch for RCNN and DeepViral. We truncated all longer sequences than 2,000 amino acids to only the first 2,000 for RCNN, due to the maximum sequence length limit of the model (similarly, first 1,000 amino acids for DeepViral). RCNN and DeepViral (seq) were implemented and evaluated for the entirety of the test set. For DeepViral variants with feature embeddings, a limited number of protein pairs, i.e. 2% of the test set, do not have relevant features available (some proteins are obsolete due to database updates) and thus are excluded from the test set. We evaluated both RCNN and the variants of DeepViral for 5 times and report the mean of the metrics.
